# Selectivity of the time-dependent *M. tuberculosis* LeuRS inhibitor ganfeborole is driven by target vulnerability

**DOI:** 10.1101/2025.07.31.667927

**Authors:** Mingqian Wang, YongLe He, Siobhan A. Cohen, Amanda R. Strohm, Gauri Shetye, Scott G. Franzblau, Stephen G. Walker, Dickon Alley, Peter J. Tonge

**Affiliations:** Center for Advanced Study of Drug Action, and Department of Chemistry, Stony Brook University, Stony Brook, NY 11794-3400, United States; Institute for Tuberculosis Research, College of Pharmacy, University of Illinois at Chicago, 833 South Wood Street, Chicago, IL 60612, United States; Department of Oral Biology and Pathology, Stony Brook University, Stony Brook, NY, 11794, United States; AN2 Therapeutics, Menlo Park, CA 94027, United States; Department of Biomedical Genetics, University of Rochester, Rochester, NY 14642, United States

**Keywords:** ganfeborole, leucyl-tRNA-synthetase, post-antibiotic effect, *Mycobacterium tuberculosis*, residence time

## Abstract

Ganfeborole (GSK3036656) inhibits the *Mycobacterium tuberculosis* leucyl-tRNA-synthetase (mtLeuRS) and is in phase 2a clinical trials for the treatment of tuberculosis. Here we show that ganfeborole is a time-dependent inhibitor of mtLeuRS (IC_50_ 1 nM) and generates a post-antibiotic effect of 77 h at 50xMIC (MIC 0.058 μM) with *M. tuberculosis* H37Rv (*Mtb*), indicating that mtLeuRS is a highly vulnerable drug target and supporting the excellent in vivo efficacy of the drug. Ganfeborole is also a potent time-dependent inhibitor of *Escherichia coli* LeuRS (ecLeuRS, IC_50_ 2 nM), however no antibacterial activity is observed towards *E. coli* up to 1 mM ganfeborole despite the observation that less potent ganfeborole analogs have antibacterial activity. To rationalize this observation, we propose that ganfeborole forms a complex with AMP that binds to the ecLeuRS editing site but does not impact aminoacylation. In support, addition of 12.5 μM norvaline generates a ganfeborole MIC of 0.4 μM since ecLeuRS is unable to hydrolyze norvaline-tRNA^Leu^. Additionally, mutations that reduce the affinity and residence time of ganfeborole-AMP on ecLeuRS result in antibacterial activity. We propose that the activity of ganfeborole towards *Mtb* is because mtLeuRS is a highly vulnerable target so that only low levels of enzyme need to be inhibited by the ganfeborole-tRNA^Leu^ complex in contrast to ecLeuRS, which we previously demonstrated is a low vulnerability target.

---

New drugs are needed to treat tuberculosis (TB), which is the leading cause of death from a single infectious agent.^1^ Drug-sensitive TB is treated by a cocktail of 2-4 drugs for 6-9 months, and either noncompliance and/or deviation from this treatment regime is a common cause of drug resistance.^2^ However, only three new drugs, bedaquiline, delamanid, and pretomanid, have been approved in the last decade for the treatment of drug-resistant TB, emphasizing the importance of developing new classes of antibiotics with novel mechanisms of action.^3,4^

The benzoxaborole ganfeborole (GSK3036656, **Figure 1**) recently entered Phase 2a clinical trials for the treatment of TB infection.^5^ Ganfeborole inhibits leucyl-tRNA synthetase (LeuRS) through the oxaborole tRNA trapping (OBORT) mechanism,^6^ in which the inhibitor forms a covalent adduct with the ribose diol of the terminal adenosine nucleotide (Ade76) of tRNA^Leu^, and traps the tRNA in the LeuRS editing site thereby inhibiting esterification of tRNA^Leu^ by Leu (**Figure 1**). Oxaborole inhibitors have also been discovered for LeuRS enzymes from other pathogens, including the first-in-class antifungal tavaborole (AN2690)^7,8^ and epetraborole (AN3365), which has potent activity toward Gram-negative bacteria.^9^ Ganfeborole was identified through an extensive med chem campaign which demonstrated the importance of the halogen at the 4 position (Cl in the case of ganfeborole) and the 2-hydroxyethoxy group at the 7 position.^10,11^ This study demonstrated that ganfeborole was highly selective for the *M. tuberculosis* LeuRS (mtLeuRS) compared to the human homologs, with reported IC_50_ values of 0.2 μM, 132 μM and >300 μM for mtLeuRS, human cytoplasmic LeuRS and human mitochondrial LeuRS, respectively.^11^

**Figure 1.**
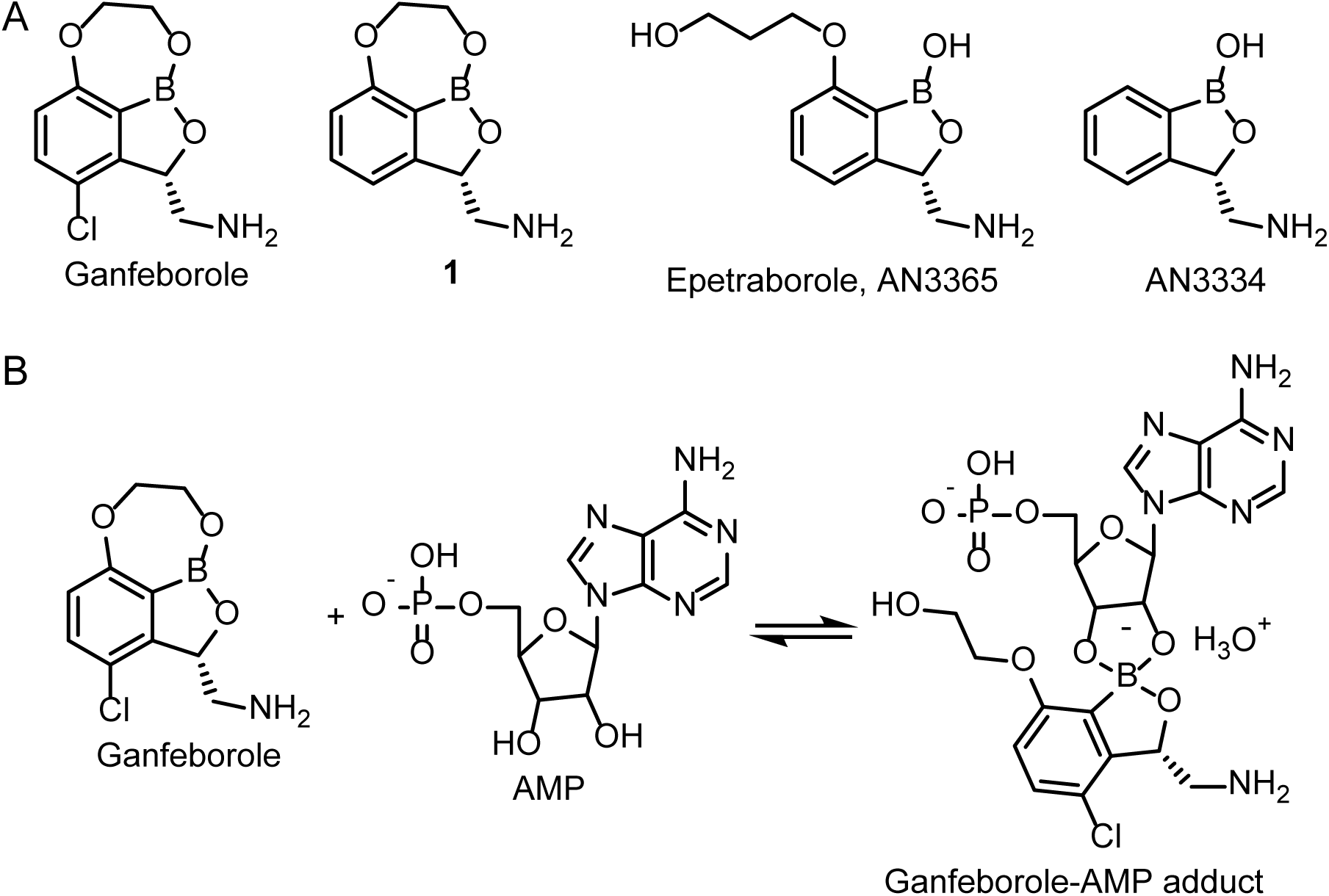
Benzoxaborole structures and the reaction of ganfeborole with AMP. (A) Structures of ganfeborole, **1**, epetraborole and AN3334. (B) Reaction of ganfeborole with AMP.

Benzoxaboroles are time-dependent inhibitors of LeuRS,^9,12^ and we initially set out to compare and contrast the binding kinetics of ganfeborole and analog **1**, which lacks the 4-Cl group, with the LeuRS enzymes from *M. tuberculosis* (mtLeuRS) and *Escherichia coli* (ecLeuRS). In the course of this work, we discovered that ganfeborole and **1** are potent inhibitors of both LeuRS enzymes but that only **1** demonstrates antibacterial activity toward *E. coli*. We subsequently demonstrate that the addition of 12.5 μM norvaline (Nva) results in a >1000-fold decrease in the MIC of ganfeborole for *E. coli,* supporting a mechanism in which a complex of ganfeborole with AMP blocks the ecLeuRS editing site without inhibiting aminoacylation. Inhibition of the ecLeuRS editing function only becomes critical in the presence of significant quantities of mischarged tRNA^Leu^, which cannot be hydrolyzed on the editing site.

## RESULTS

The benzoxaboroles are time-dependent inhibitors of LeuRS.^9,12^ We first quantified the inhibition of ecLeuRS and mtLeuRS by ganfeborole and compound **1**, an analog lacking the 4-Cl group. *E. coli* and *M. tuberculosis* tRNA^Leu^ were prepared by *in vitro* transcription, and in each case, a decrease in IC_50_ value was observed after 1 h preincubation of inhibitor, enzyme, and tRNA^Leu^ consistent with slow-onset inhibition (**Figure 2**, **Table 1**). The final IC_50_ values for ganfeborole and **1** with ecLeuRS were 2 and 5 nM, respectively, while those for mtLeuRS were 1 and 2 nM, respectively.

**Figure 2.**
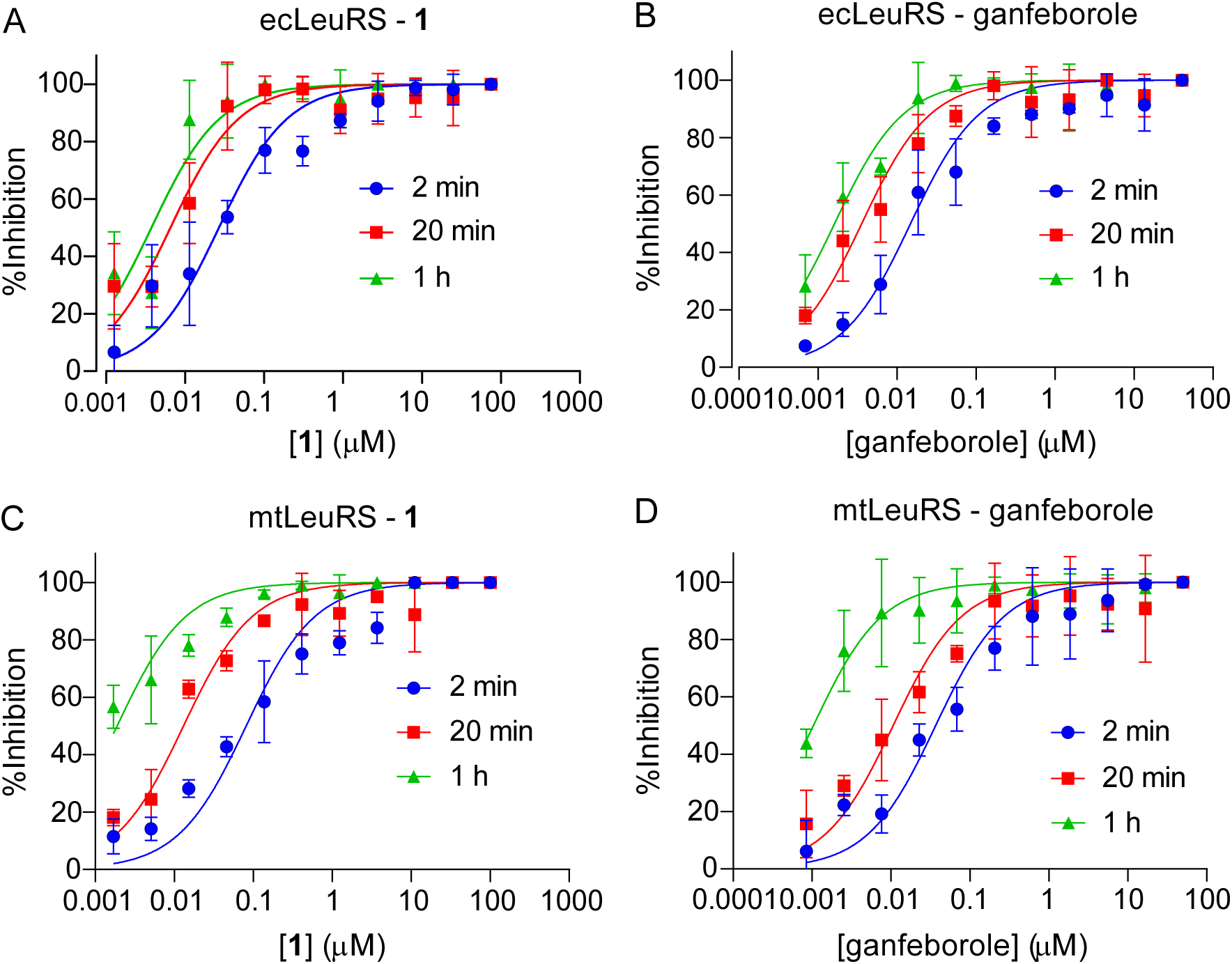
Concentration-response plots for the time-dependent inhibition of LeuRS. (A) ecLeuRS and **1**, (B) ecLeuRS and ganfeborole, (C) mtLeuRS and **1**, and (D) mtLeuRS and ganfeborole. Compound **1** and ganfeborole were preincubated with 800 pM ecLeuRS and 20 μM ectRNA^Leu^ or 800 pM mtLeuRS and 20 μM mttRNA^Leu^ for 2 min, 20 min and 1 h. The reaction was initiated with 10 μM ATP and 20 μM L-Leu, and then quenched with 20 mM EDTA after 5 min. Fluorescence polarization (FP) was converted to [AMP] using an AMP/ATP FP standard curve, and the inhibition% was calculated and normalized using the DMSO-only control. The data was fit to a concentration-response equation using GraphPad Prism in which the Hill slope was constrained to 1.0. Each experiment was performed in at least three independent biological replicates, with the error bars representing the standard deviation from the mean. Each experiment was performed in at least two independent biological replicates and the errors are the standard deviations from the mean.

**Table 1.**
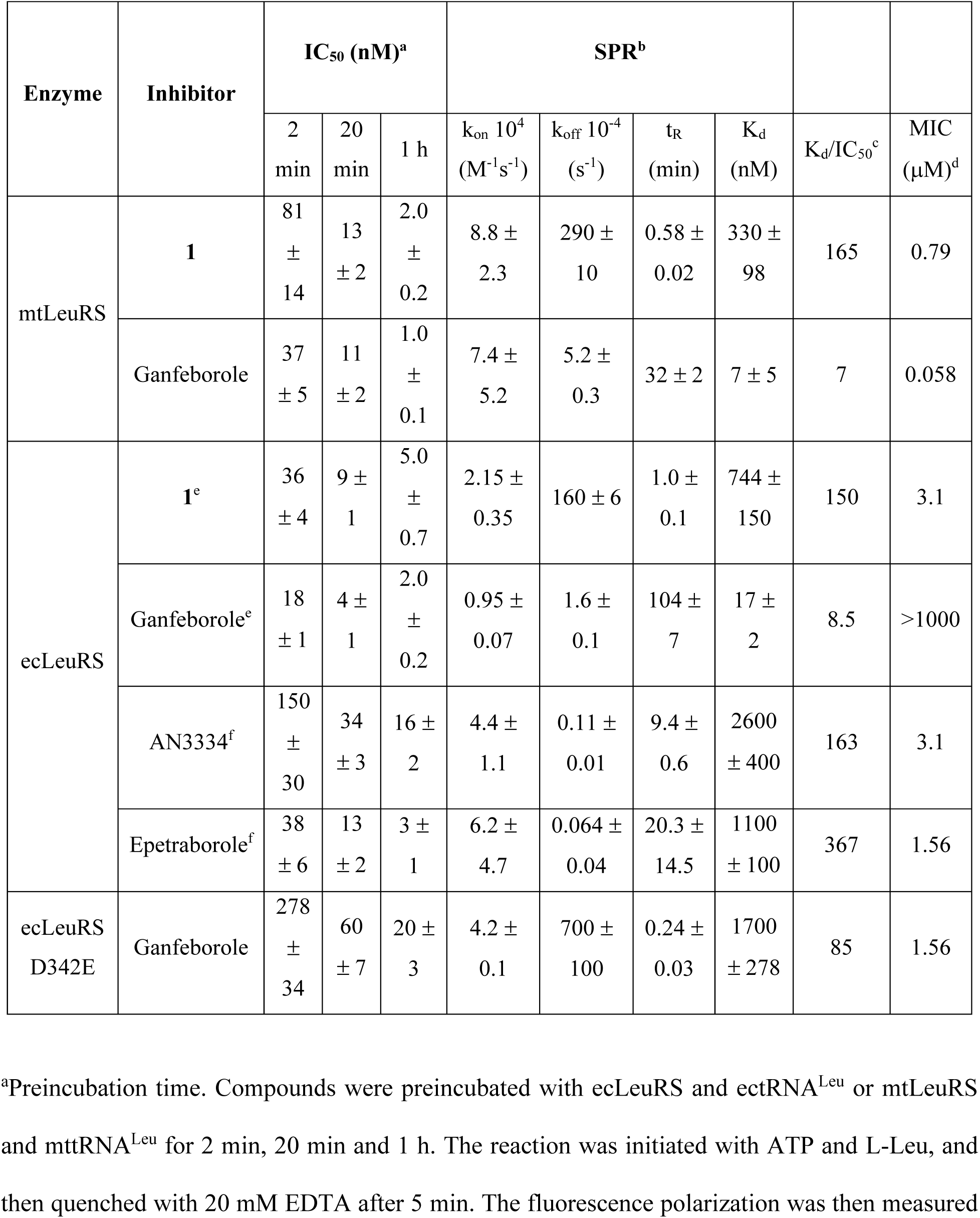
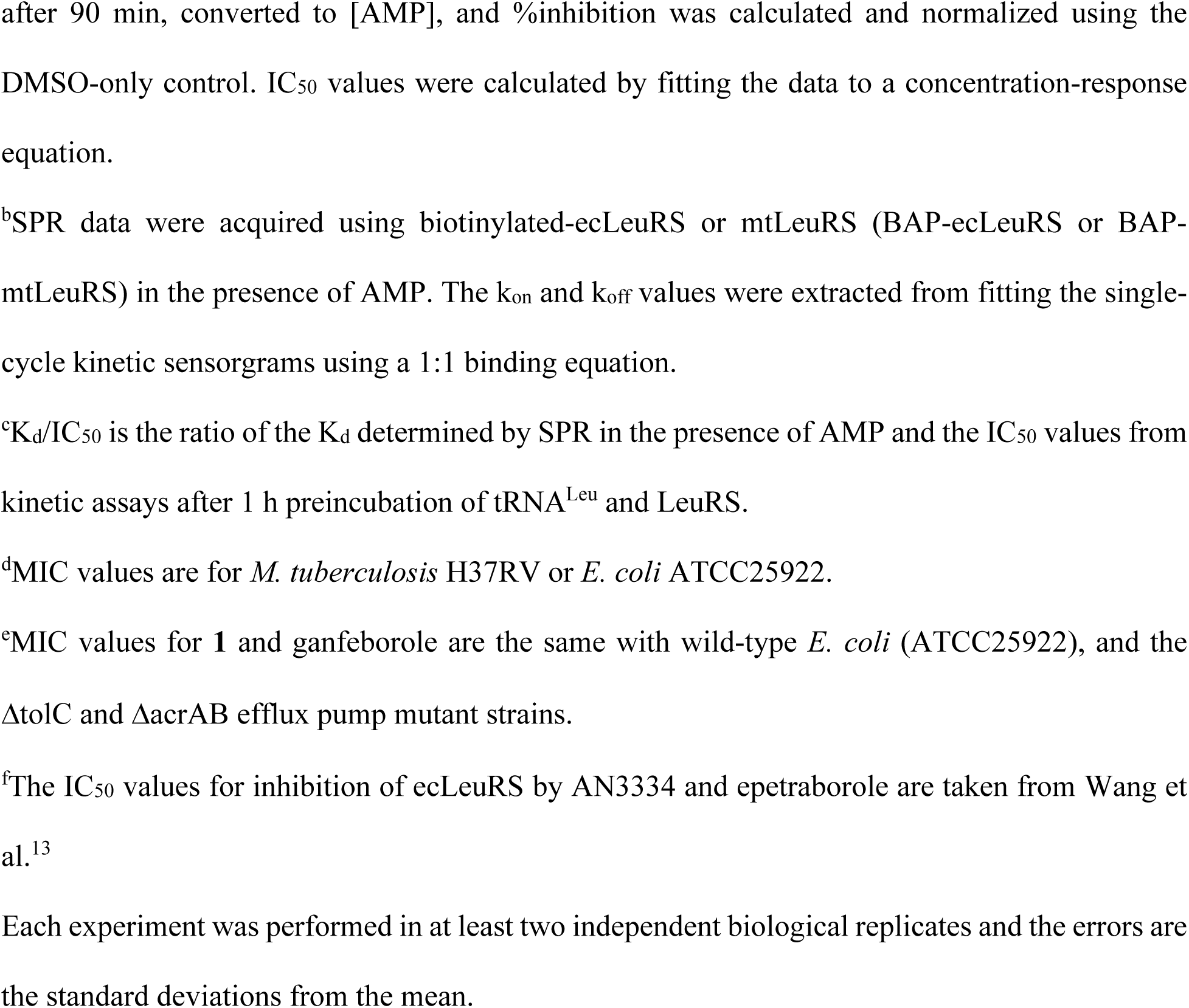
Binding kinetics and microbiological activity of the LeuRS inhibitors.

| Enzyme | Inhibitor | IC<sub>50</sub> (nM)<sup>a</sup> |  |  | SPR<sup>b</sup> |  |  |  |  |  |
| --- | --- | --- | --- | --- | --- | --- | --- | --- | --- | --- |
|  |  | 2 min | 20 min | 1 h | k<sub>on</sub> 10<sup>4</sup> (M<sup>-1</sup>s<sup>-1</sup>) | k<sub>off</sub> 10<sup>-4</sup> (s<sup>-1</sup>) | t<sub>R</sub> (min) | K<sub>d</sub> (nM) | K<sub>d</sub>/IC<sub>50</sub><sup>c</sup> | MIC (μM)<sup>d</sup> |
| mtLeuRS | **1** | 81 ± 14 | 13 ± 2 | 2.0 ± 0.2 | 8.8 ± 2.3 | 290 ± 10 | 0.58 ± 0.02 | 330 ± 98 | 165 | 0.79 |
|  | Ganfeborole | 37 ± 5 | 11 ± 2 | 1.0 ± 0.1 | 7.4 ± 5.2 | 5.2 ± 0.3 | 32 ± 2 | 7 ± 5 | 7 | 0.058 |
| ecLeuRS | **1<sup>e</sup>** | 36 ± 4 | 9 ± 1 | 5.0 ± 0.7 | 2.15 ± 0.35 | 160 ± 6 | 1.0 ± 0.1 | 744 ± 150 | 150 | 3.1 |
|  | Ganfeborole<sup>e</sup> | 18 ± 1 | 4 ± 1 | 2.0 ± 0.2 | 0.95 ± 0.07 | 1.6 ± 0.1 | 104 ± 7 | 17 ± 2 | 8.5 | >1000 |
|  | AN3334<sup>f</sup> | 150 ± 30 | 34 ± 3 | 16 ± 2 | 4.4 ± 1.1 | 0.11 ± 0.01 | 9.4 ± 0.6 | 2600 ± 400 | 163 | 3.1 |
|  | Epetraborole<sup>f</sup> | 38 ± 6 | 13 ± 2 | 3 ± 1 | 6.2 ± 4.7 | 0.064 ± 0.04 | 20.3 ± 14.5 | 1100 ± 100 | 367 | 1.56 |
| ecLeuRS D342E | Ganfeborole | 278 ± 34 | 60 ± 7 | 20 ± 3 | 4.2 ± 0.1 | 700 ± 100 | 0.24 ± 0.03 | 1700 ± 278 | 85 | 1.56 |

<sup>a</sup>Preincubation time. Compounds were preincubated with ecLeuRS and ectRNA<sup>Leu</sup> or mtLeuRS and mttRNA<sup>Leu</sup> for 2 min, 20 min and 1 h. The reaction was initiated with ATP and L-Leu, and then quenched with 20 mM EDTA after 5 min. The fluorescence polarization was then measured
after 90 min, converted to [AMP], and %inhibition was calculated and normalized using the DMSO-only control. IC<sub>50</sub> values were calculated by fitting the data to a concentration-response equation.
<sup>b</sup>SPR data were acquired using biotinylated-ecLeuRS or mtLeuRS (BAP-ecLeuRS or BAP-mtLeuRS) in the presence of AMP. The $k_{on}$ and $k_{off}$ values were extracted from fitting the single-cycle kinetic sensorgrams using a 1:1 binding equation.
<sup>c</sup>K<sub>d</sub>/IC<sub>50</sub> is the ratio of the K<sub>d</sub> determined by SPR in the presence of AMP and the IC<sub>50</sub> values from kinetic assays after 1 h preincubation of tRNA<sup>Leu</sup> and LeuRS.
<sup>d</sup>MIC values are for *M. tuberculosis* H37RV or *E. coli* ATCC25922.
<sup>e</sup>MIC values for **1** and ganfeborole are the same with wild-type *E. coli* (ATCC25922), and the ΔtolC and ΔacrAB efflux pump mutant strains.
<sup>f</sup>The IC<sub>50</sub> values for inhibition of ecLeuRS by AN3334 and epetraborole are taken from Wang et al.<sup>13</sup>
Each experiment was performed in at least two independent biological replicates and the errors are the standard deviations from the mean.

The microbiological activity of the compounds against *M. tuberculosis* H37Rv and *E. coli* ATCC 25922 was assessed by determining the minimum inhibitory concentration (MIC) and post-antibiotic effect (PAE). The MIC values for **1** and ganfeborole with H37Rv were 0.79 and 0.058 μM, respectively (**Table 2**). PAE values were then determined by first exposing bacterial cultures to 1x, 10x, 30x, and 50xMIC of each inhibitor for 2.5 h, followed by 10,000-fold dilution into fresh media, after which the rate of bacterial growth was quantified.^14,15^ The PAE values were concentration-dependent, increasing to 42 and 77 h with 50xMIC exposure for **1** and ganfeborole, respectively (**Figure 3**, **Table 2**). As controls, PAE values were also obtained for isoniazid and rifampicin. In agreement with previous studies, isoniazid generated no detectable PAE at any concentration, whereas exposure to rifampicin resulted in a concentration dependent PAE that increased to 72 h at 50xMIC.^15^ The PAE values observed for **1** and ganfeborole support the conclusion that mtLeuRS is a highly vulnerable drug target,^13,16–19^ in agreement with gene titration experiments using CRISPRi.^20^

**Table 2.**
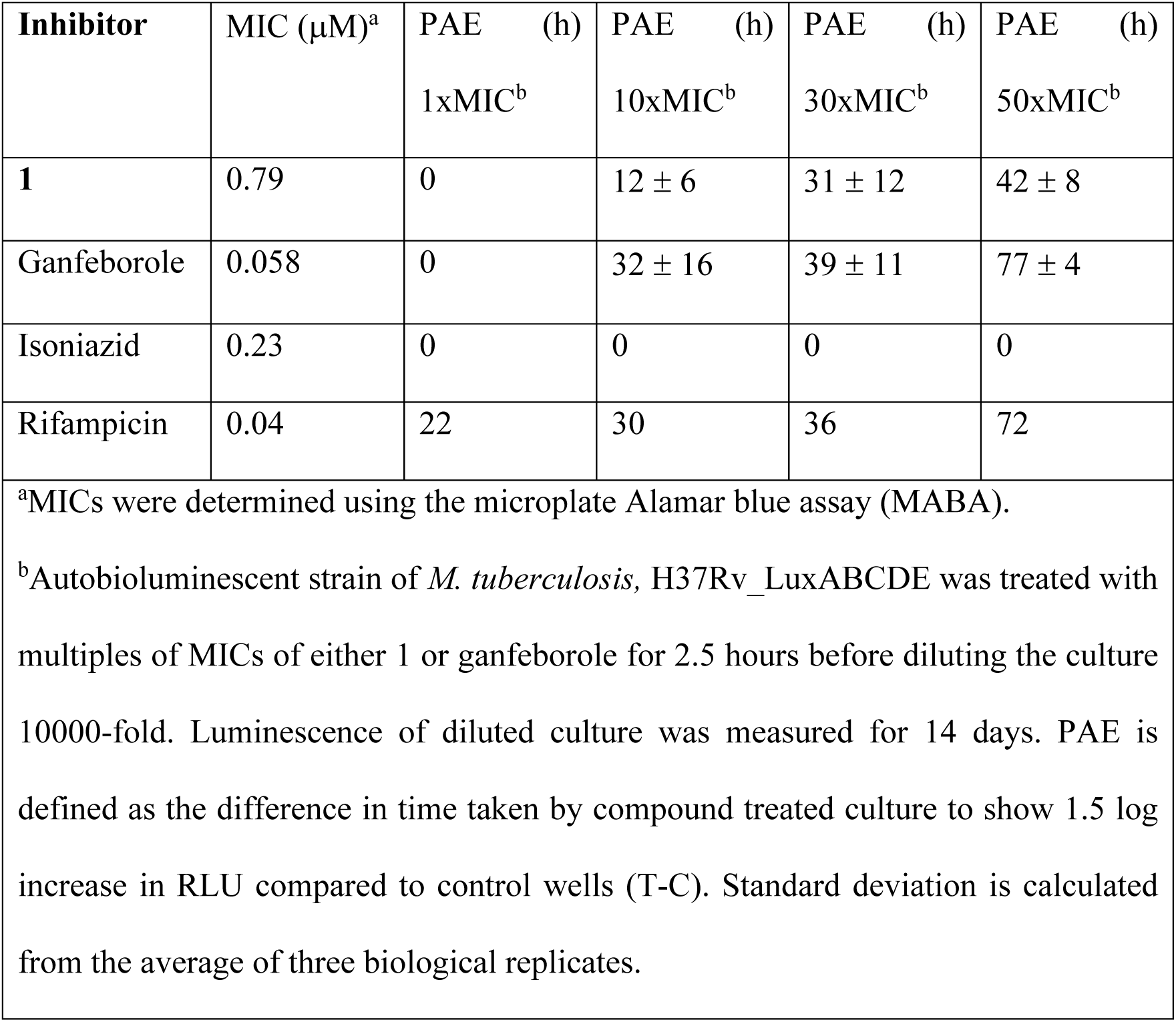
MIC and PAE values of 1 and ganfeborole with *M. tuberculosis* H37Rv.

**Figure 3.**
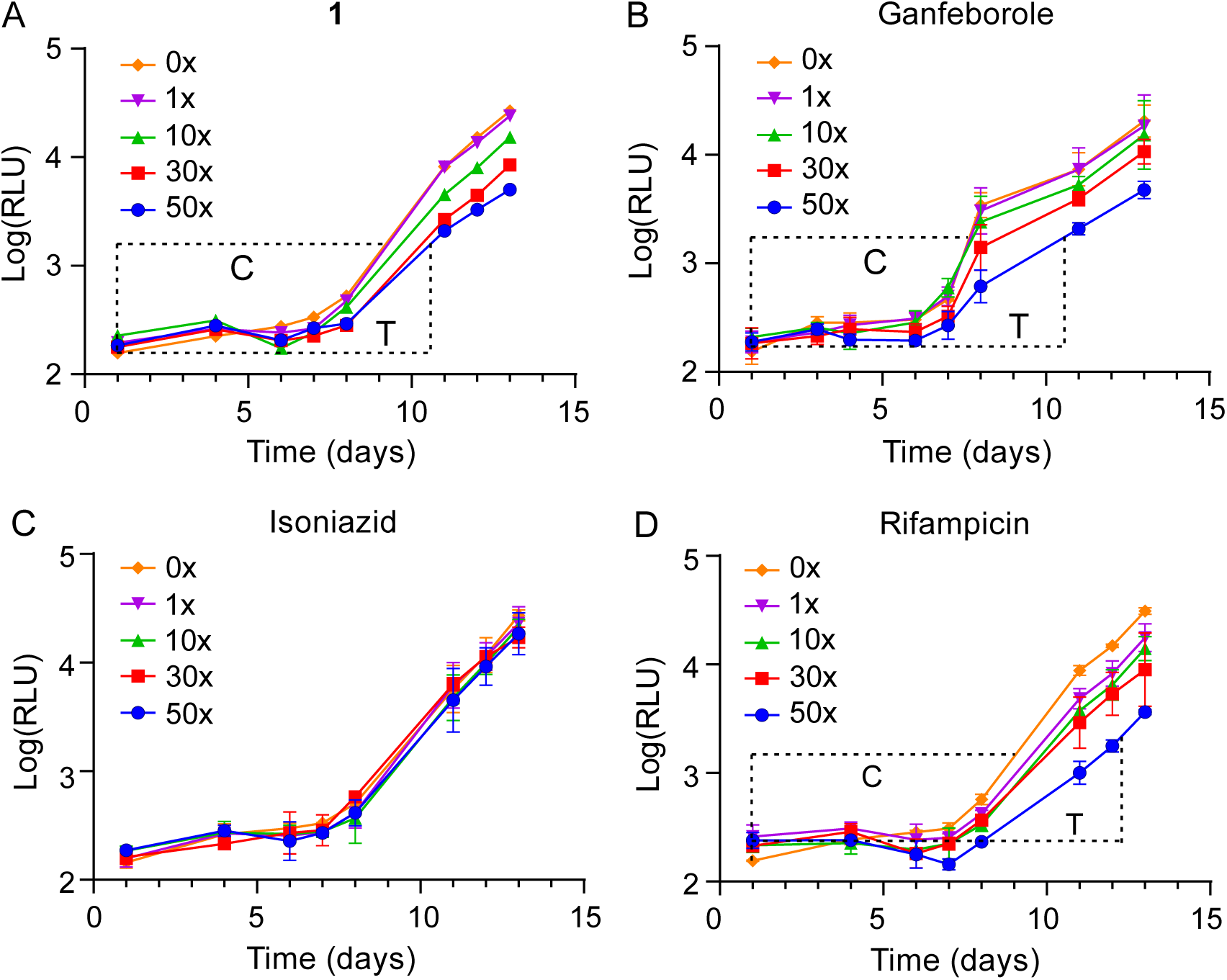
PAE of 1, ganfeborole, isoniazid and rifampicin with *M. tuberculosis* H37Rv. PAE of (A) **1**, (B) ganfeborole, (C) isoniazid and (D) rifampicin at 0, 1xMIC, 10xMIC, 30xMIC and 50xMIC. An autobioluminescent strain of *M. tuberculosis,* H37Rv_LuxABCDE, was treated with multiples of MICs of compounds for 2.5 h before diluting the culture 10,000-fold. The luminescence of the diluted culture (RLU) was measured for 14 days. The PAE is defined as the difference in time taken by compound treated culture to show 1.5 log increase in RLU compared to control wells (T-C). Dashed lines show the PAE induced by 50xMIC of compound. Error bars represent the standard deviation calculated from the average of three biological replicates.

For *E. coli*, the MIC of **1** was 3.1 μM. However, ganfeborole had no detectable antibacterial activity up to 1 mM (**Table 1**). This result was surprising given that other close structural analogs of ganfeborole have antibacterial activity against *E coli*. For instance, AN3334 and epetraborole have MIC values of 3.1 and 1.56 μM (**Table 1**), and DS86760016 has an MIC of 1 μM.^9,21^ Ganfeborole also had no antibacterial activity in the *E. coli ΔtolC* and *ΔacrAB* strains (**Table 1**), suggesting that efflux did not account for the lack of activity. We then demonstrated that ganfeborole could effectively antagonize the antibacterial activity of **1** in the wild-type strain, where increasing concentrations of ganfeborole raised the MIC of **1** from 3.1 μM to 6.25 μM at 0.1 μM ganfeborole, and 200 μM at 6.25 μM ganfeborole (**Table 3**). This indicates that ganfeborole is able to penetrate the bacteria and displace **1** from ecLeuRS; in other words, ganfeborole can bind to ecLeuRS but not inhibit the synthesis of Leu-tRNA^Leu^.

**Table 3:** MIC values of the benzoxaboroles in combination and in the presence of Nva and Leu.

| Compound | MIC (μM) *E. coli* ATCC25922<sup>a</sup> |
| --- | --- |
| **1** | 3.1 |
| **1** + 12.5 μM Nva | 3.1 |
| **1** + 0.2 μM ganfeborole | 6.25 |
| **1** + 6.25 μM ganfeborole | 200 |
| Ganfeborole | >1000 |
| Ganfeborole + 12.5 μM Nva | 0.4 |
| Ganfeborole + 100 μM Nva + 10 μM Leu | 1.56 |
| Ganfeborole + 100 μM Nva + 100 μM Leu | 1600 |
| <sup>a</sup>*E. coli* ATCC25922 with serial dilutions of combinations of **1**, ganfeborole, Nva and Leu in 96-well plates containing M9 media. MIC was determined with the lowest drug concentration where no visible growth was observed.<br>Each experiment was performed in at least three independent biological replicates. |  |

It had previously been reported that AMP can replace tRNA^Leu^ and that the benzoxaborole-AMP complex is a potent inhibitor of LeuRS.^22^ We speculated that the ganfeborole-AMP complex occupies the editing site of ecLeuRS and does not prevent aminoacylation from occurring. To evaluate this hypothesis, we repeated the antibacterial assays under conditions where the editing function of ecLeuRS would become more important. This was accomplished by adding norvaline (Nva) to the media, given that tRNA^Leu^ misloaded with Nva is known to be removed by LeuRS.^23,24^ In support of the hypothesis, whereas Nva had no effect on bacterial growth up to 200 μM and did not affect the MIC of **1**, the addition of 12.5 μM Nva resulted in a ganfeborole MIC of 0.4 μM, which is a more than a 1000-fold increase in antibacterial activity (**Table 3**). The synergistic activity of Nva and ganfeborole was confirmed using a checkboard assay which generated a FICI value << 0.5 (**Figure 4**). In addition, time-kill assays demonstrated that the combination of ganfeborole and Nva resulted in cidality (**Figure 5**). In agreement with the proposed mechanism, it was subsequently shown that Leu could antagonize the effect of Nva on the antibacterial activity of ganfeborole. For instance, the MIC of Nva with 6.4 μM ganfeborole increased to 25 μM in the presence of 3.1 μM Leu, and to 1.6 mM with 200 μM Leu (**Table 3**, **Figure 4** and **5**).

**Figure 4.**
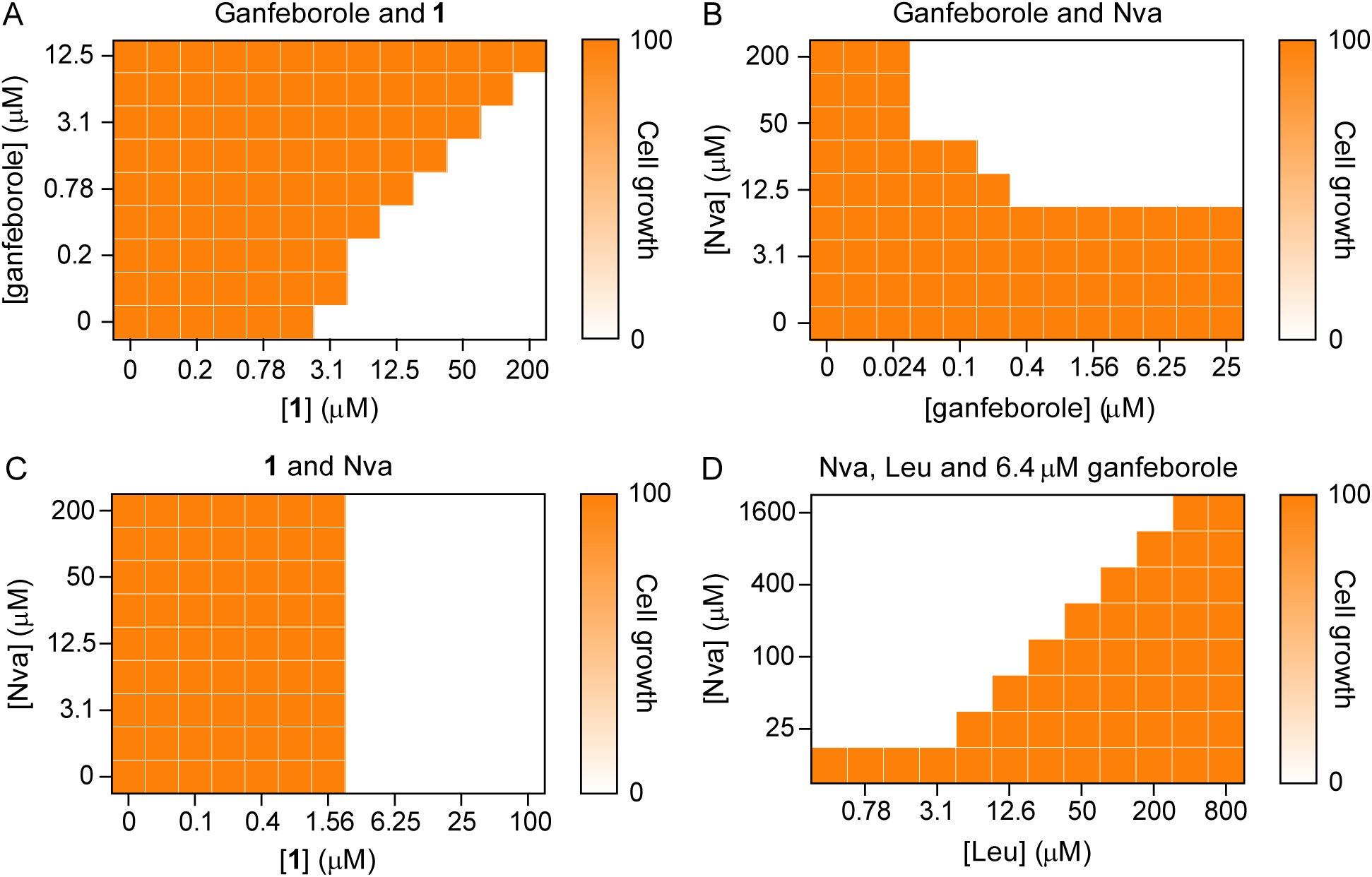
Checkerboard MIC assay of 1 and ganfeborole with Nva and Leu in *E. coli* ATCC25922. (A) Ganfeborole and **1**, (B) ganfeborole and Nva, (C) **1** and Nva, and (D) Nva and Leu with 6.4 µM ganfeborole. Two-fold serial dilutions of the two compounds were added to the 96-well plate that contained 5x10^5^ CFU/mL *E. coli* cells/well in M9 media. Visual turbidity in each well was determined after 16 h incubation at 37°C. Each experiment was performed in at least three independent biological replicates.

**Figure 5.**
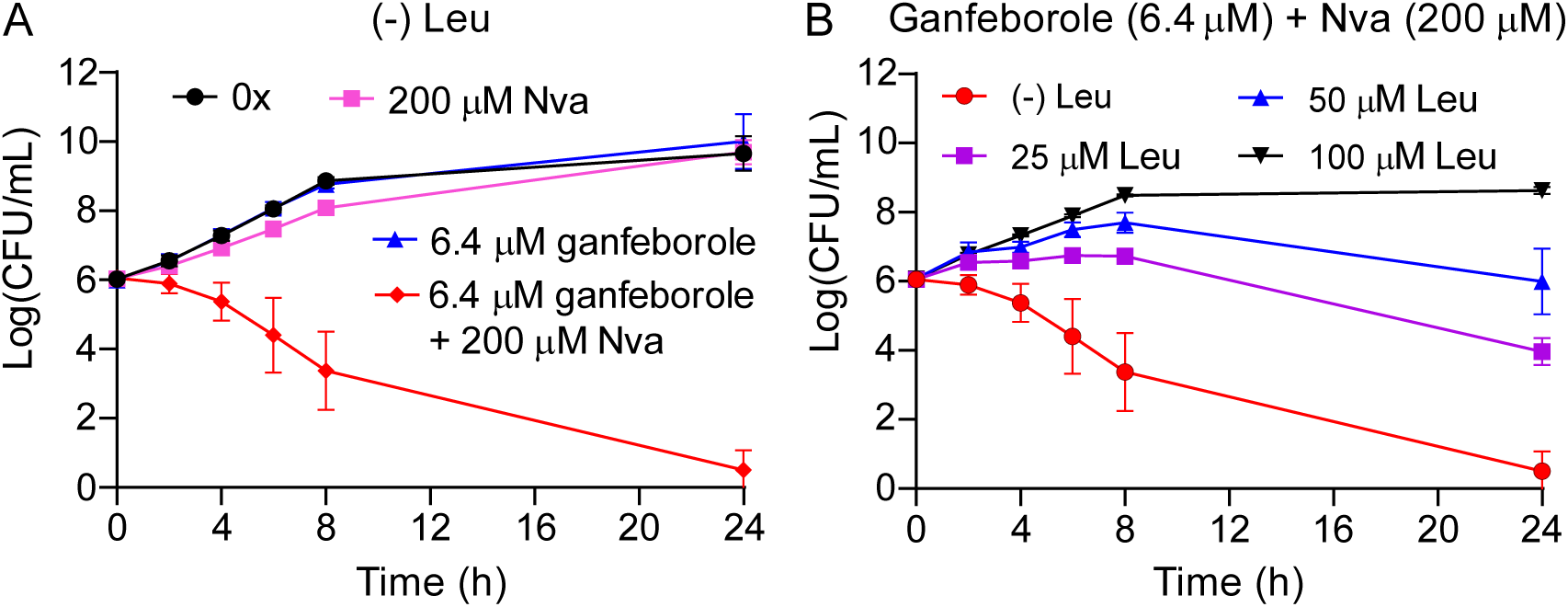
Time-kill assay of 1 and ganfeborole with Nva and Leu against *E. coli* ATCC25922. Time-kill experiments were performed with (A) ganfeborole, Nva with 0 μM Leu. (B) 6.4 µM ganfeborole and 200 µM Nva with 0, 25, 50, and 100 µM Leu. Experiments were performed in the M9 medium. Aliquots were taken every hour from the culture to record the growth of *E. coli* over time. Each experiment was performed in at least three independent biological replicates, with the error bars representing the standard deviation from the mean.

The checkerboard and time-kill experiments demonstrate that the activity of ganfeborole is impacted by Nva, which we speculated was due to the inability of the ecLeuRS editing site to correct the misloading of tRNA^Leu^ by Nva. To support this hypothesis, we generated strains of *E. coli* that were resistant to **1** by exposing *E. coli* to 10xMIC (31 µM) of **1**. Whole genome sequencing of four strains that had MIC values of >250 μM for **1** revealed mutations at D342E, G333D, D345N, and V335D in the ecLeuRS editing site. In contrast to wild-type *E. coli*, ganfeborole has MIC values of 1.56 µM – 6.25 µM with the four mutant strains, which indicates that ecLeuRS is the target of ganfeborole and that the inhibitor binds in the editing domain (**Table 4**).

**Table 4.** MIC values of ganfeborole in strains of *E. coli* that are resistant to 1.

| *E. coli* mutant strains<sup>a</sup> | MIC of **1** (μM) | MIC of ganfeborole (μM)<sup>b</sup> |
| --- | --- | --- |
| Wild-type | 3.1 | >1000 |
| D342E | >1000 | 1.56 |
| G333D | 250 | 6.25 |
| D345N | >1000 | 3.1 |
| V335D | 500 | 6.25 |

<sup>a</sup>*E. coli* mutant strains resistant to **1** were generated by treating 10<sup>8</sup> cells with 31 μM **1**. Single colonies were picked after overnight growth on Mueller-Hinton agar. *E. coli* mutant strains were treated with 2-fold dilutions of ganfeborole in 96-well plates in the CAMH medium. The MIC was defined as the lowest drug concentration where no visible growth was observed. Each experiment was performed in at least three independent biological replicates.

To further characterize the binding of ganfeborole and AMP to LeuRS we used surface plasmon resonance (SPR). For both enzymes, ganfeborole bound ∼30-40-fold more potently to LeuRS than compound **1**, with K_d_ values of 7 and 17 nM for mtLeuRS and ecLeuRS, respectively (**Figure 6** and **Table 1**). In addition, in keeping with the slow-onset inhibition observed from the time-dependent shift in IC_50_ values, ganfeborole was found to have a long residence time on both enzymes with t_R_ values of 100 and 32 min for ecLeuRS and mtLeuRS, respectively (**Table 1**).

**Figure 6.**
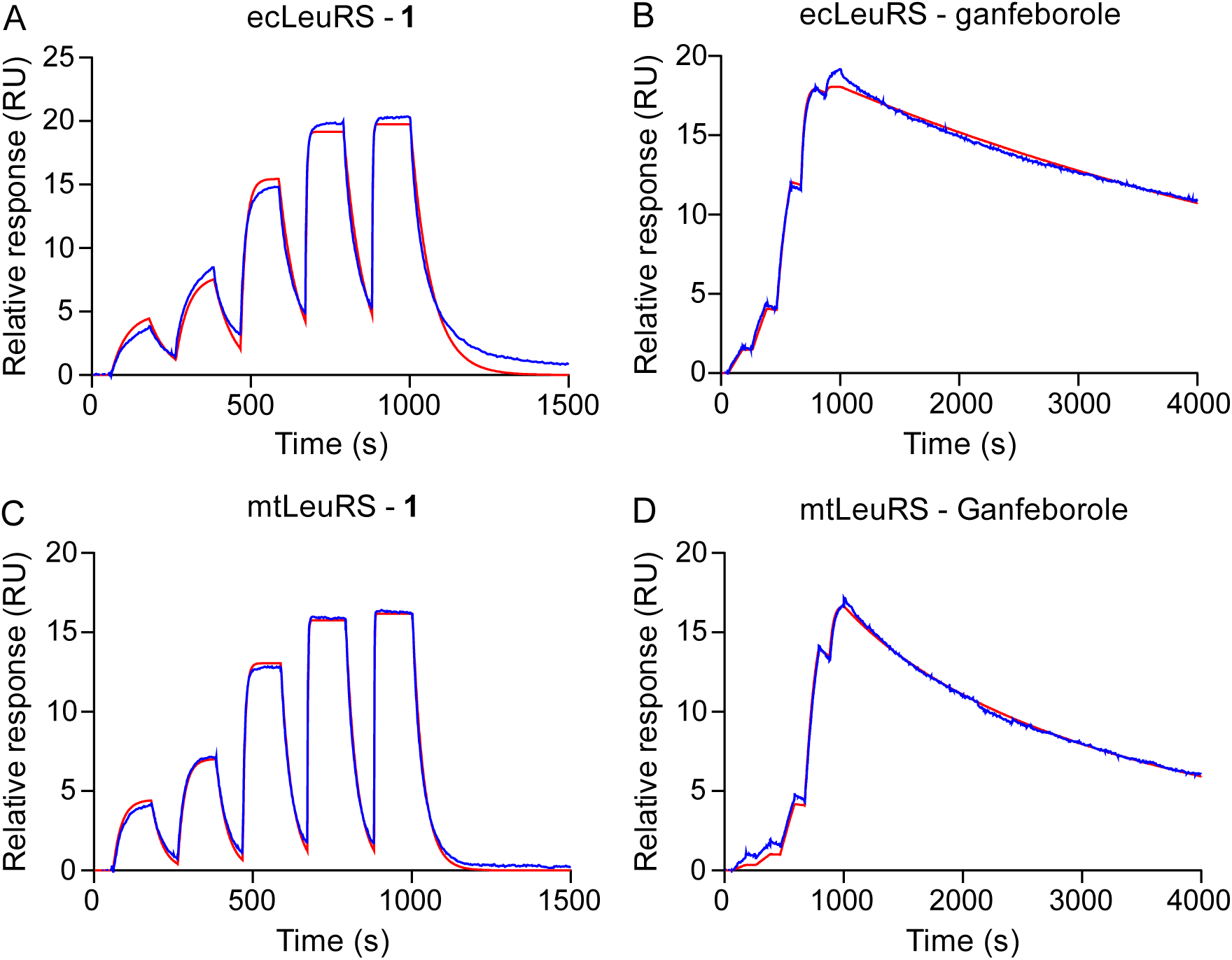
SPR sensorgrams for the binding of 1 and ganfeborole to LeuRS in the presence of AMP. Single-cycle kinetic binding sensorgrams of (A) ecLeuRS and **1**, (B) ecLeuRS and ganfeborole, (C) mtLeuRS and **1**, and (D) mtLeuRS and ganfeborole. BAP-ecLeuRS or BAP-mtLeuRS was immobilized on the streptavidin sensor chip. Five different concentrations of **1** and ganfeborole were preincubated with AMP and then coinjected onto the SPR. The data was fitted to a 1:1 binding model using the Biacore T200 Evaluation Software to extract all the binding parameters. Each experiment was performed in at least two independent biological replicates.

As the D342E mutation showed the greatest MIC shift for ganfeborole, we then analyzed the impact of this mutation on the binding kinetics of ganfeborole and **1** (**Figure 7** and **Table 1**). Whereas ganfeborole had an IC_50_ of 2 nM for ecLeuRS following 1 h preincubation, this value increased 10-fold to 20 nM for the D342E mutant. In addition, using SPR, it was shown that ganfeborole-AMP became a rapid reversible inhibitor of D342E ecLeuRS and that the K_d_ increased from 17 nM to 1.7 µM.

**Figure 7.**
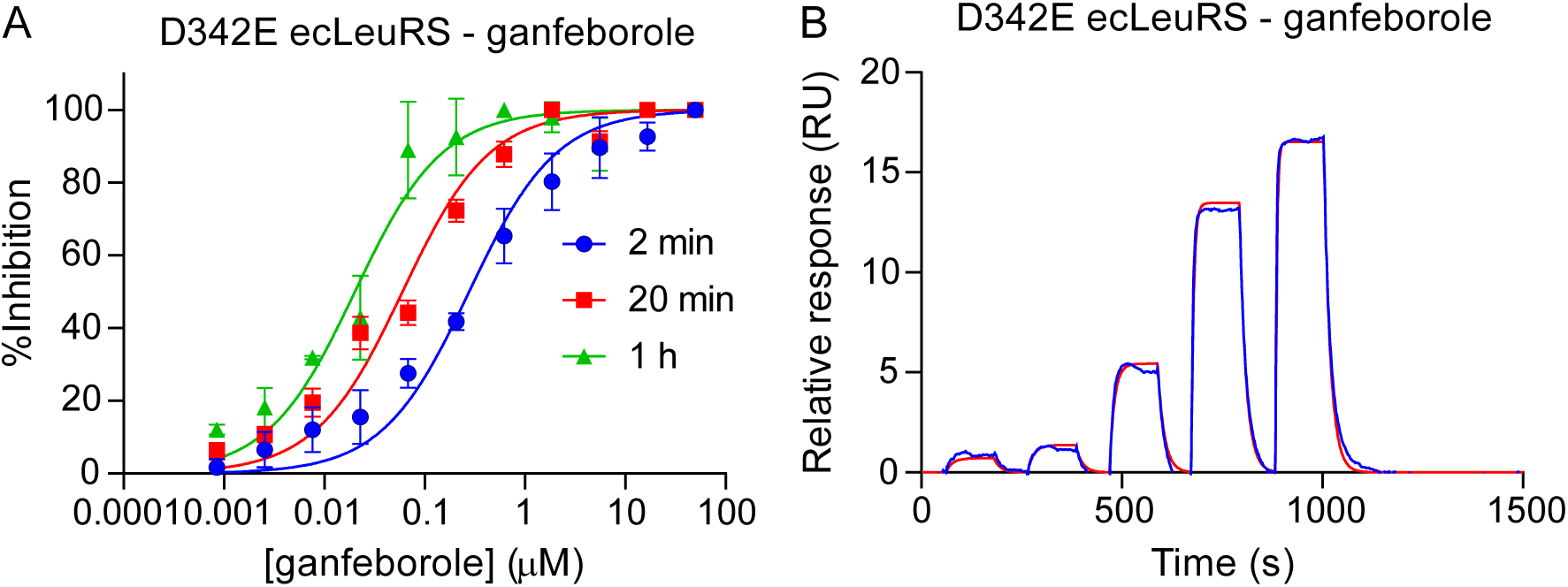
Binding of ganfeborole to D342E ecLeuRS. (A) Time-dependent inhibition of ganfeborole to D342E ecLeuRS. Ganfeborole was preincubated with 800 pM D342E ecLeuRS and 20 μM ectRNA^Leu^ for 2 min, 20 min and 1 h. The reaction was initiated with 10 μM ATP and 20 μM L-Leu, and then quenched with 20 mM EDTA after 5 min. Fluorescence polarization was converted to [AMP] using a standard curve, and the %inhibition was calculated and normalized using the DMSO-only control. The data was fit to a concentration-response equation using GraphPad Prism in which the Hill slope was constrained to 1. (B) Single-cycle kinetic binding sensorgrams of D342E ecLeuRS with ganfeborole. BAP-D342E ecLeuRS was immobilized on the streptavidin sensor chip. Five different concentrations of ganfeborole were preincubated with AMP and then coinjected onto the SPR chip. The data was fit to a 1:1 binding model using the Biacore T200 Evaluation Software to extract the binding parameters.

## DISCUSSION

Ganfeborole (GSK3036656) is in clinical trials for the treatment of TB.^5^ This compound has potent antibacterial activity toward drug-sensitive and drug-resistant strains of *M. tuberculosis* and has an MIC of 0.058 μM against *M. tuberculosis* H37Rv. Initial characterization of ganfeborole demonstrated >500-fold selectivity compared to human LeuRS homologs, with a reported IC_50_ of 0.2 μM for mtLeuRS with 20 min preincubation.^11^ The benzoxaboroles are time-dependent inhibitors of LeuRS with reported residence times of 1-8 h for AN3334 and 23 h for AN3213 (the racemate of epetraborole, AN3365) with ecLeuRS and tRNA^Leu^.^9,12^ However, the time-dependent inhibition of mtLeuRS had not been previously studied. To compare and contrast the binding kinetics of ganfeborole with those of other benzoxaborole LeuRS inhibitors, we have studied the time-dependent activity of ganfeborole with mtLeuRS and the homolog from *E. coli*.

Using an activity assay, we showed that the IC_50_ values for inhibition of mtLeuRS by ganfeborole and the structural analog **1** that lacks the 4-Cl substituent decrease by more than 10-fold to 1-2 nM following a 1 h preincubation of enzyme, inhibitor, and mttRNA^Leu^. We then used SPR to further analyze the binding kinetics of the two compounds with mtLeuRS. Using AMP as a surrogate for tRNA^Leu^,^22^ we showed that ganfeborole binds ∼50-fold more potently to mtLeuRS than **1** and has a residence time on the enzyme of 32 min compared to <1 min for **1**. These data thus confirm that ganfeborole is a slow-onset inhibitor of mtLeuRS, and the previously reported IC_50_ of 0.2 μM likely reflects the initial rapid binding of the inhibitor to the enzyme in a two-step binding mechanism.

We previously demonstrated that drug-target residence time can correlate with the post-antibiotic effect (PAE).^17–19,25,26^ which is the delay in bacterial regrowth following exposure and removal of an antibiotic.^27^ The PAE is an important component of antibacterial dosing regimens,^19,25^ consequently, we determined the PAE generated by ganfeborole and **1** for H37Rv and observed PAE values of 77 h and 42 h, respectively, following 50xMIC exposure. The PAEs are similar to those generated by protein synthesis inhibitors such as rifampicin (68-77 h) and streptomycin (32 h),^28,29^ and the ClpC1 inhibitor ecumicin (80 h),^15^ but considerably longer than those caused by other TB drugs such as moxifloxacin (0-0.3 h), ethambutol (2 h) and isoniazid (0-20 h).^15,28,29^ The PAEs generated by TB drugs become considerably longer when used in combination; for instance, isoniazid and rifampicin have a PAE of 160 h,^28,29^ and thus, the ability of ganfeborole to elicit a PAE indicates that mtLeuRS is a highly vulnerable target and may prove to be an important aspect of human dosing regimens.

Ganfeborole and **1** were found to have similar biochemical activity with ecLeuRS, the LeuRS homolog in *E. coli*. Both compounds were time-dependent inhibitors with IC_50_ values of 2-5 nM following 1 h preincubation and K_d_ values of 17 and 744 nM, respectively, determined by SPR in the presence of AMP. In addition, the ganfeborole-AMP complex had a residence time of 104 min on ecLeuRS compared to 1 min for the complex of AMP with **1**. However, whereas **1** had an MIC of 3.1 μM against a wild-type strain of *E. coli*, ganfeborole has no antibacterial activity up to 1 mM against either wild-type or pump mutant strains of *E. coli*. The lack of antibacterial activity was not due to an inability of ganfeborole to penetrate *E. coli*, since ganfeborole antagonized the activity of **1**. We subsequently showed that addition of Nva resulted in potent ganfeborole antibacterial activity which we speculate is due to increased dependence on the editing function of ecLeuRS due to misloading of ectRNA^Leu^ by Nva. In agreement, addition of Leu antagonized the effect of Nva on the ganfeborole MIC.

Oxaboroles inhibit aminoacyl-tRNA synthetases by trapping uncharged tRNA in the editing site through the formation of a complex with the terminal A76 ribose. However, studies by Hoffman et al. revealed that oxaboroles can also bind to LeuRS in the presence of nucleotides such as AMP (**Figure 8**).^22^ In the case of ecLeuRS, the results with ganfeborole demonstrate that the oxaborole inhibition complex formed with AMP does not inhibit the formation of leucyl-tRNA^Leu^ and only leads to antibacterial activity when the LeuRS editing function is needed. In contrast, **1** and other oxaboroles that have antibacterial activity against *E. coli* in the absence of Nva must preferentially form a complex with tRNA^Leu^ on the enzyme. Given that the concentration of nucleotides such as AMP and ATP is at least 10-fold higher than tRNA^Leu^ (1-5 mM compared to 100-200 μM),^22,30^ the inhibition complex formed with tRNA must be normally 10-100-fold more potent than that formed with (e.g.) AMP, which is reasonable given that tRNA will form many more interactions with the enzyme. Indeed, comparison of the IC_50_ and K_d_ values obtained from activity assays in the presence of tRNA^Leu^ and SPR using AMP, respectively, shows that the K_d_/IC_50_ ratio for ecLeuRS is ≥150-fold for **1**, AN3334 and epetraborole. However, the ratio drops to 8.5 for ganfeborole indicating the 5’ chloro group has preferentially stabilized the enzyme-inhibitor complex form with AMP compared to tRNA^Leu^. In the D342E editing site mutant the K_d_/IC_50_ ratio for ganfeborole has increased to 85-fold, explaining the antibacterial activity of ganfeborole toward the mutant strain.

**Figure 8:**
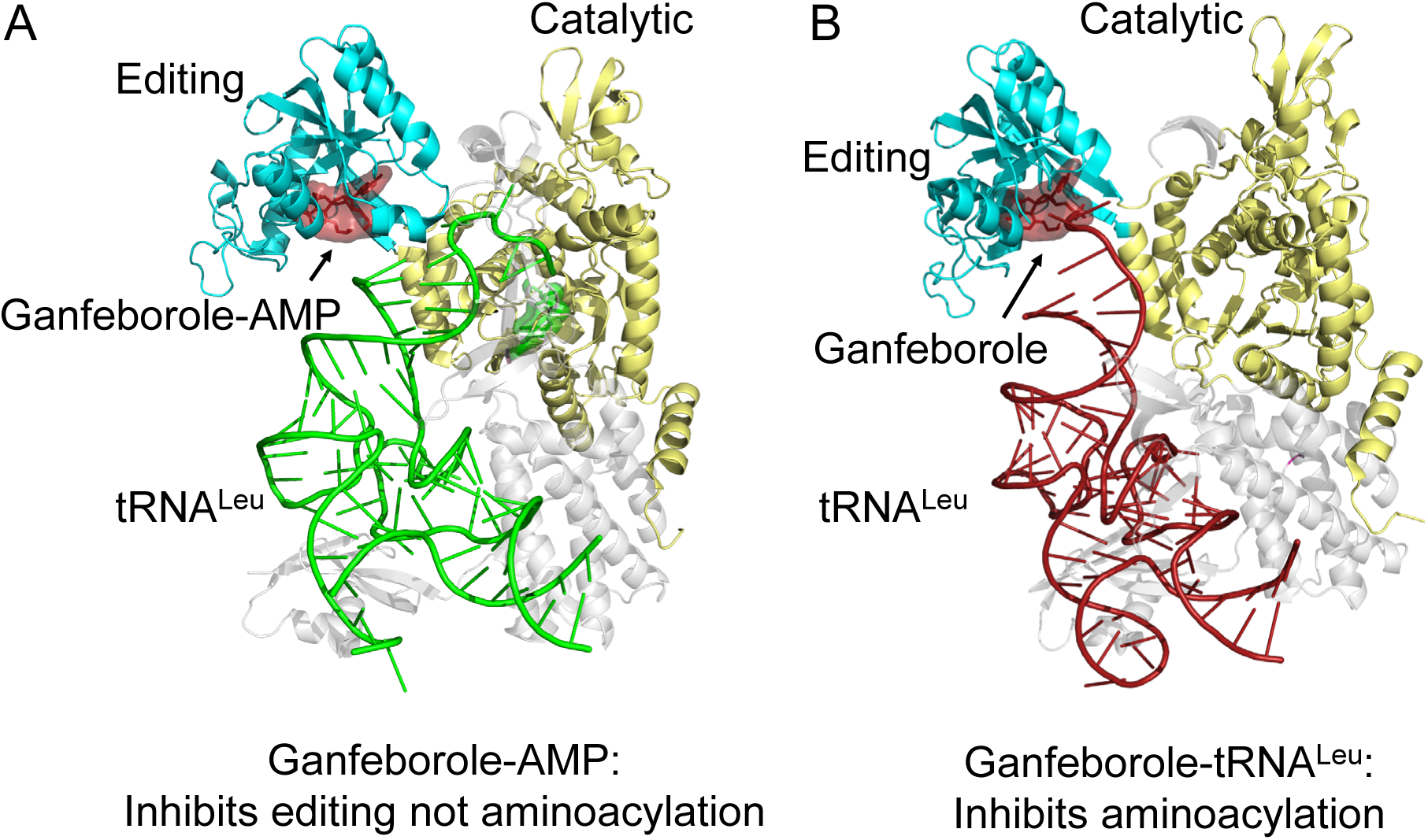
Oxaboroles Bind to LeuRS either in complex with tRNA^Leu^ or AMP. (A) The ganfeborole-AMP complex binds to the editing site of LeuRS, inhibiting editing but not aminoacylation. (B) The ganfeborole-tRNA^Leu^ complex inhibits aminoacylation. The relative amounts of each complex depend on the concentrations of LeuRS, ganfeborole, tRNA^Leu^ and nucleotides such as AMP. The ganfeborole-AMP complex is significantly more potent than the complex formed with other oxaboroles, thus the relative ratio of complex A to B is significantly higher for ganfeborole. Since mtLeuRS is more vulnerable than ecLeuRS, the required fractional occupancy of mtLeuRS by ganfeborole-tRNA^Leu^ is smaller than that in ecLeuRS.

In contrast to *E. coli*, ganfeborole has potent activity toward *M. tuberculosis* despite similar differences in the K_d_/IC_50_ ratios for **1** and ganfeborole (160 and 7-fold, respectively) compared to ecLeuRS. Here we demonstrate that mtLeuRS is a highly vulnerable target in contrast to ecLeuRS which we previously showed was a low vulnerability target.^13^ Thus, one possible explanation for the differential activity of ganfeborole in the two organisms is that a smaller fraction of mtLeuRS compared to ecLeuRS has to be occupied by ganfeborole-tRNA^Leu^ to inhibit bacterial growth (**Figure 8**).^25^ In addition, while ganfeborole has a significant residence time on mtLeuRS with AMP, the value of 32 min is short relative to the ∼24 h doubling time of *M. tuberculosis*, whereas it is long (104 min) on ecLeuRS compared to the ∼20 min doubling time of *E. coli*. In either case, the high vulnerability of mtLeuRS and ability of ganfeborole to generate a significant post-antibiotic effect in *Mtb*, support the excellent in vivo efficacy of ganfeborole in the recent successfully completed Phase 2a early bactericidal activity clinical trial.

## MATERIALS AND METHODS

### Compounds and Bacterial Strains

Ganfeborole, epetraborole, AN3334 and **1** were gifts from AN2 Therapeutics. L-Norvaline and L-leucine were purchased from Fisher Scientific. Adenosine 5’-monophosphate monohydrate was purchased from Sigma-Aldrich. *E. coli* ATCC 25922 was purchased from ATCC. *E. coli* ΔacrAB was kindly provided by Prof. Vincent Tam at the University of Houston, and *E. coli* ΔtolC was kindly provided by Prof. Zgurskaya at the University of Oklahoma. Muller Hinton Broth (Cation-Adjusted) and Muller Hinton Agar were purchased from BD Difco.

### Cloning, Expression, and Purification of LeuRS

The *E. coli* leucyl tRNA synthetase was amplified from *E. coli* ATCC 25922 by PCR using primer

1 ATGCAAGAGCAATACCGCCCGGAAGAGATAGAATC and primer 2 TTAGCCAACGACCAGATT GAGGAGTTTACCGGGTAC. The gene insert was gel extracted and cloned into the pEMB54 vector using the Gibson Assembly method, which is an ampicillin-resistant, arabinose-inducible vector with a pMB1 origin of replication and 6XHis-Smt3 under the PBAD (araBp) promoter, including a multiple cloning site containing BamHI and HindIII sites following the Smt3 sequence. Genes cloned into pEMB54 via BamHI/HindIII are fused in-frame with 6XHis-Smt3, leaving a serine linker following the C-terminal double glycine of Smt3. The codon-optimized Leucyl tRNA synthetase from *Mycobacterium tuberculosis* (mtLeuRS, WP_128882958.1) was sub-cloned into pET28TEV vector via NdeI/BamHI, which is fused in-frame with the N-terminal 6XHis Tag followed by a TEV cleavage site. The constructs utilized for the SPR study were inserted with a C-terminal Biotin Acceptor Peptide (BAP) using primer 1 GCGCAGAAAATTGAATGGCATGAAAAGCTTGGCTGTTTTG and primer 2 TTCAAAAATATCGTTCAGGCCACCCACAACCAGGTTCAGC for ecLeuRS and primer 1 GCAGAAAATTGAATGGCATGAATAAGGATCCGAATTCGAG and primer 2 GCTTCAAAAATATCGTTCAGGCCAATCACCAGGTTAACCAG for mtLeuRS. The amplified products were circulated with the Kinase, Ligase, and DpnI reaction mix (KLD Enzyme Mix, NEB). The final ligation was transformed into chemically competent DH5α cells, and the construct was mini-prepped from a colony, after which the sequence was confirmed by Sanger DNA sequencing.

The *E. coli* Bl21 (DE3) component cells were transformed with the pEMB54 ecLeuRS or pET28TEV mtLeuRS construct by heat shock at 42°C and plated on LB-Agar plate containing 100 μg/mL of ampicillin (ecLeuRS) or 50 μg/mL of kanamycin (mtLeuRS). The overnight culture was prepared by inoculating LB Miller broth, containing appropriate antibiotic, with a single colony and incubating at 37°C in an orbital shaker (250 rpm). After overnight incubation at 37°C, 10 mL of the overnight culture was inoculated into 500 mL of 2XYT media broth with appropriate antibiotic. The cultures were incubated at 37°C in an orbital shaker (250 rpm) until the OD600 ∼0.6 - 0.8 was obtained, and then, the temperature was lowered to 18°C. Once the OD600 was ∼0.8 - 1, 0.1% of arabinose (Sigma-Aldrich) or 1 mM of isopropyl β-D-1-thiogalactopyranoside (IPTG, Gold Biosciences) was added to induce protein expression, and the culture was incubated with shaking for an additional 16 h. Cells from the 1 L culture were harvested by centrifugation at 5000 rpm (4°C) for 20 min, and the cell pellet was stored at −20°C.

After thawing, the pellet was resuspended in 35 mL of lysis buffer (25 mM Tris-HCl pH 8, 150 mM NaCl, 2 mM MgCl_2_, and 5 mM imidazole, buffer A) and lysed by sonication. The cell debris was removed by ultracentrifugation at 40,000 rpm for 1 h (4°C) and filtered by a 40 μm constrainer.

The filtered supernatant was then loaded into a 5 mL Ni-NTA column (GE) previously equilibrated with buffer A. After all the supernatant was applied to the column, the column was then washed with 10-20 CV of buffer A containing 10-30 mM of Imidazole. A gradient of 30-500 mM of Imidazole elution was performed, and the protein was eluted at 500 mM Imidazole. Fractions containing protein were pooled and concentrated by a 10 kDa cut-off concentrator and desalted using the HiPrep 26/10 desalting column (GE) with buffer A. Fractions containing protein were pooled, and 250 μg of homemade UlpI SUMO protease (ecLeuRS) or TEV protease (mtLeuRS) was added to cleave the SUMO tag/6XHis tag at 4°C overnight. The cleaved protein was loaded onto the Ni-NTA column and washed 8 CV with buffer A. Fraction containing LeuRS was concentrated by a 10 kDa cut-off concentrator and loaded onto a HiLoad 16/600 Superdex-75 size exclusion column previously equilibrated with the store buffer (25 mM Tris-HCl pH 8, 150 mM NaCl, 2 mM DTT, 10% glycerol). For the sample that contains the BAP, biotinylation was performed prior to size exclusion chronography. The BAP-protein was desalted into the biotinylation buffer (50 mM Bicine pH 8, 30 mM NaCl), followed by adding 10 mM ATP, 10 mM magnesium acetate, and 1 mM d-Biotin. ∼300 μg of homemade BirA was added into the sample and incubated at 25°C for 1 to 2 h, and the BirA was separated using size exclusion chromatography. Pure protein fractions were collected, and the purity of the protein was confirmed by SDS-PAGE. The concentration of the protein was determined by Absorbance spectrum using extinction coefficient at 280 nm (ε280 = 169,765 M^-1^cm^-1^ for ecLeuRS, 189,330 M^-1^cm^-1^ for mtLeuRS).

### *In Vitro* Transcription of tRNA^Leu^

All oligodeoxynucleotides used as templates for enzymatic transcription were obtained from Integrated DNA Technologies. The oligonucleotides used for construction of the *E. coli* tRNA^Leu^ gene were PO_4_- GGATCC**TAATACGACTCACTATA**GCGGGAGTGGCGAAATTGGTAGACG<u>CACCAGATTTAG</u> and TGGTGCGGGAGGCGAGACTTGAACTCGCACACCTTGCGGCGCCAGAAC<u>CTAAATCTGGTG</u>. Oligonucleotides were designed such that the sense strand corresponding to the 5’ end of the sequence possessed a 12 bp overlap with the 3’ antisense strand. The underlined portions of the sequence represent the overlapping regions, and the bold text indicates the T7 RNA polymerase promoter. Oligonucleotides were mixed to an equimolar concentration of 4 μM in a reaction solution containing 400 μM dNTPs, 10 mM Tris-HCl, pH 7.5, 10 mM MgSO_4_, 7.5 mM DTT, and 50 U/mL Klenow fragment polymerase (New England Biolabs).^31^ The mixture was cycled between 10°C and 37°C at 30 s intervals for eight cycles. The PCR products were extracted from a 3% agarose gel by cutting the band with the correct molecular weight. The gel-extracted gene was then amplified by PCR using the Phusion High-Fidelity DNA Polymerase (Thermo Fisher Scientific) with primers 5’-GGATCCTAATACGACTCACTATAGCGGGAG and 5’TGGTGCGGGAGGCGA GACTTG. The second PCR amplified product was purified by the PCR clean-up kit (Qiagen). The oligonucleotides used for the construction of the *M. tuberculosis* tRNA^Leu^ gene were PO_4_-GGATCC**TAATACGACTCACTATA**GGGCGAGTGGCGGAATGGCAGACGCGCTG GCTTCAGGTGCCAGTGTCCTTCGGGACGTGGGGGTTCAAGTCCCCCTTCGCCCACCA (Synthesized by GenScript).

The *in vitro* transcription was performed using the HiScribe T7 High Yield RNA Synthesis Kit (NEB). Briefly, 7.5 mM of NTP, 10 mM of T7 RNA polymerase mix, 0.75x of reaction buffer, and 300 ng of purified DNA template were used for the *in vitro* transcription reaction. The reaction mixture was incubated at 37°C for 16 h, followed by the addition of 2 U DNase digest for 15 min at 37°C. Subsequently, 0.1 volume of 3 M sodium acetate was added to the final tRNA^Leu^ product, and the tRNA^Leu^ was precipitated using ethanol.

### IC_50_ Measurements using LeuRS Aminoacylation-Activity Assay

The aminoacylation reaction was performed in 50 mM HEPES buffer pH 8, containing 1 mM MgCl_2_ and 1 mM DTT. The formation of the AMP product was monitored over time using the Transcreener AMP^2^ /GMP^2^ FP Assay Kit (BellBrook). ecLeuRS or mtLeuRS (800 pM) was preincubated with 20 µM ectRNA^Leu^ or mttRNA^Leu^ and 2-fold serial dilutions of the ganfeborole or **1** from 2 min to 1 h, after which the reaction was initiated by adding 10 µM ATP and 20 µM L-leucine. After 5 min at 25°C, the reaction was quenched by adding the detection mixture, which included 20 mM EDTA. After 90 min at 25°C, the fluorescence polarization was read on a plate reader (BioTek Synergy Neo2) with 620 nm excitation and 680 nm emission. The enzyme activity was determined by converting the mP value to the AMP concentration using the standard curve. The data were fitted to a nonlinear regression model using GraphPad Prism to determine the IC_50_ value for each compound.

### Residence Time Measurements using Surface Plasmon Resonance

The kinetic assay was performed by SPR using a Biacore T200 (Cytiva). BAP-ecLeuRS full length or BAP-mtLeuRS full length was immobilized on the Serie S SA Sensor chip (Cytiva) at ∼3500 response units after the chip surface was rinsed with a mixed solution of 1 M NaCl and 50 mM NaOH, and tested for binding at 25°C. One of the two flow paths was left empty as a reference for nonspecific binding. The running buffer comprises 10 mM HEPES, 150 mM NaCl, and 0.5% (v/v) Surfectant P20, pH 7.4 (HBS-P buffer, Cytiva). The ganfeborole or **1** stock solution was diluted into the running buffer containing 1% DMSO and 1 mM AMP to yield a series of working stocks, which included five different dilution concentrations for single-cycle kinetics, and preincubated at RT for 1 h before injections. The flow rate was 60 µL/min, and the injection time was set to 120 s with a 60 s equilibration time between injections and a 3600 s dissociation time. Data was referenced using a reference flow cell in line with the active flow cell. All data were analyzed using the Biacore T200 Evaluation Software (GE Healthcare). All experiments were done in at least duplicates.

### Minimum Inhibitory Concentration

1. *E. coli* ATCC 25922 was grown to mid-log phase (OD600 of 0.5 - 0.7) in M9 minimal media, consisting of M9 salts (6.78 g/L Na_2_HPO_4_, 3 g/L KH_2_PO_4_, 0.5 g/L NaCl, 1 g/L NH_4_Cl), 5 g/L glucose, 1 mM MgSO_4_, 0.1 mM CaCl_2_, 100 μL trace metal solution, 33 μM thiamine at 37°C in an orbital shaker. An inoculum of 10^6^ CFU/mL was added to each well, containing 2-fold dilutions of inhibitors in a round-bottom 96-well plate. The MIC is defined as the minimum concentration of the inhibitor at which no visible growth showed after 16 h at 37°C. All experiments were performed in triplicate.

Minimum inhibitory concentration against replicating *Mtb* is determined using a previously described Microplate Alamar Blue Assay (MABA).^14^ Briefly, compound stocks were prepared in dimethyl sulfoxide (DMSO) at 100x of the highest desired final concentration. Subsequently, 2 µL of the compound DMSO stock was transferred to the assay plate containing 100 µL of Middlebrook 7H12 media (4.7 g 7H9 broth, containing 1 g casitone (Bacto), 5 g bovine serum albumin (BSA), 4 mg catalase and 5.6 mg palmitic acid for 1 L media). Two-fold serial dilutions of the compounds was performed in triplicates in the assay plate. Plates are then inoculated with *Mtb* strain H37Rv (ATCC 27294) to achieve a final density of ∼ 1 × 10^5^ CFU/mL in each well and incubated for 7 days at 37°C. At the end of 7 days, a resazurin dye/tween 80 mixture (0.6 mM resazurin dye and 12 µL of 20% Tween 80) was added to each well, and the plates were further incubated for an additional 18 to 24 h at 37°C. Fluorescence was measured using a CLARIOstar (BMG LABTECH, Ortenberg, Germany) plate reader on day 8. The MIC is defined as the lowest concentration of compound that resulted in a 90%reduction in fluorescence compared to the controls.

### Post-antibiotic Effect in *Mtb*

The post-antibiotic effect in *Mtb* was determined using a previously described method.^15^ Briefly, 2 µL of compound stock was transferred to 200 µL of bacterial culture of 1 × 10^6^ CFU/mL density in row B of 96-well plates. Each compound was tested at 50X, 30X, 10X, and 1X MIC concentrations. Wells from rows C to G were filled with 180 µL of 7H12 media (4.7 g 7H9 broth), containing 1 g casitone (Bacto), 5 g bovine serum albumin (BSA), 4 mg catalase and 5.6 mg palmitic acid for 1 L media). Plates were incubated in a humidified incubator for 2.5 h at 37°C, after which 20 µL of bacterial culture from row B were serially diluted 10-fold up to row G (10^5^-fold dilution), and the plate was transferred back to the incubator. Following serial dilutions, the luminescence of the plate was read every day up to day 14. Although cultures were serially diluted up to 10^5^, graphs were plotted, and PAE was calculated based on the 10^4^ dilution. PAE is defined as the difference in time taken by compound treated culture to show a 1.5 log increase in RLU compared to control wells.

### Checkerboard MIC Assay and FIC Index

*E. coli* ATCC 25922 was grown to mid-log phase (OD 600 of 0.5 - 0.7) in cation-adjusted Mueller Hinton (CAMH) media at 37°C in an orbital shaker. Then, 100 µL of 10^6^ CFU/mL cells were added to each well in a round-bottom 96-well plate containing 2-fold dilutions of two compounds. The MIC was defined as the minimum concentration of the inhibitor at which no visible growth was observed after 16 h at 37°C. All experiments were performed in triplicate.

The fractional inhibitory concentration index (FICI) was determined in order to calculate the synergistic effect between the two compounds using **Equation 1**.^32,33^

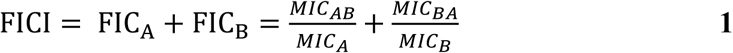

In Equation **1**, FIC_A_ and FIC_B_ are the FIC values of compounds A and B, respectively, and MIC_A_ is the MIC of compound A, MIC_B_ is the MIC of compound B, MIC_AB_ is the MIC of compound A in the presence of compound B and MIC_BA_ is the MIC of compound B in the presence of compound A. A FICI ≤ 0.5 indicates a synergistic effect, 0.5 < FICI ≤ 1 indicates an additive effect, 1 < FICI ≤ 4 indicates indifference and FICI > 4 indicates an antagonistic effect.

### Time-Kill Assays in *E. coli* ATCC25922

1. *E. coli* ATCC 25922 was grown to mid-log phase (OD600 of 0.5 - 0.7) in M9 minimal media, consisting of M9 salts (6.78 g/L Na_2_HPO_4_, 3 g/L KH_2_PO_4_, 0.5 g/L NaCl, 1 g/L NH_4_Cl), 5 g/L glucose, 1 mM MgSO_4_, 0.1 mM CaCl_2_, 100 μL trace metal solution, 33 μM thiamine at 37°C in an orbital shaker. Then, an inoculum of 10^6^ CFU/mL was added to the media containing various concentrations of ganfeborole, **1**, L-norvaline, or vehicle (DMSO). Subsequently, 100 μL aliquots were taken at 0, 2 h, 4 h, 6 h, 8 h, and 24 h and plated in serial dilutions on Muller-Hinton agar plates. After incubating at 37°C overnight, CFUs were calculated by counting the colonies on each plate.

### Generation and Sequencing of *E. coli* Strains Resistant to 1

1. *E. coli* ATCC 25922 was grown in CAMH media to OD600 = 1 at 37°C in an orbital shaker.

Subsequnetly,100 µL of *E. coli* cells were plated on Mueller-Hinton agar containing 31 µM (10xMIC) of **1**. Plates were incubated at 37°C overnight and four colonies were used to inoculate 20 mL CAMH medium containing 31 µM (10xMIC) **1** at 37°C overnight in an orbital shaker. The

1. *E. coli* cells were pelleted at 37°C for 10 min at 3900 rpm. DNA extraction and Whole Genome Sequencing were performed by Admera Health.

## AUTHOR INFORMATION

### Corresponding Author

**Peter Tonge** - Center for Advanced Study of Drug Action, and Departments of Chemistry and Radiology, John S. Toll Drive, Stony Brook University, Stony Brook, NY 11794-3400, United States; Department of Biomedical Genetics, University of Rochester, Rochester, NY 14642, United States;

**M.R.K. Alley –** AN2 Therapeutics, Menlo Park, CA 94027, United States;

### Authors

**Mingqian Wang -** Center for Advanced Study of Drug Action, and Department of Chemistry, John S. Toll Drive, Stony Brook University, Stony Brook, NY 11794-3400, United States;

**Yongle He -** Center for Advanced Study of Drug Action, and Department of Chemistry, John S. Toll Drive, Stony Brook University, Stony Brook, NY 11794-3400, United States;

**Siobhan Cohen -** Center for Advanced Study of Drug Action, and Department of Chemistry, John S. Toll Drive, Stony Brook University, Stony Brook, NY 11794-3400, United States;

**Amanda Strohm -** Center for Advanced Study of Drug Action, and Department of Chemistry, John S. Toll Drive, Stony Brook University, Stony Brook, NY 11794-3400, United States;

**Gauri Shetye** - Institute for Tuberculosis Research, College of Pharmacy, University of Illinois at Chicago, 833 South Wood Street, Chicago, IL 60612, United States;

**Scott G. Franzblau** - Institute for Tuberculosis Research, College of Pharmacy, University of Illinois at Chicago, 833 South Wood Street, Chicago, IL 60612, United States;

**Stephen G. Walker -** Department of Oral Biology and Pathology, Stony Brook University, Stony Brook, NY 11794, United States;

## ACKNOWLEDGMENTS

This study was supported by the National Institutes of Health (GM149297 to P.J.T.). Y.H. was supported by a National Institutes of Health Chemistry-Biology Interface Training Grant (GM136572), and S.A.C. was supported by a National Institutes of Health Training Grant for Scholars in Biomedical Sciences (GM148331).

## ABBREVIATIONS

LeuRS: Leucyl-tRNA-Synthetase
PAE: Post-antibiotic Effect
MIC: Minimum Inhibitory Concentration
Nva: norvaline
SPR: surface plasmon resonance.

## For Table of Contents Use Only

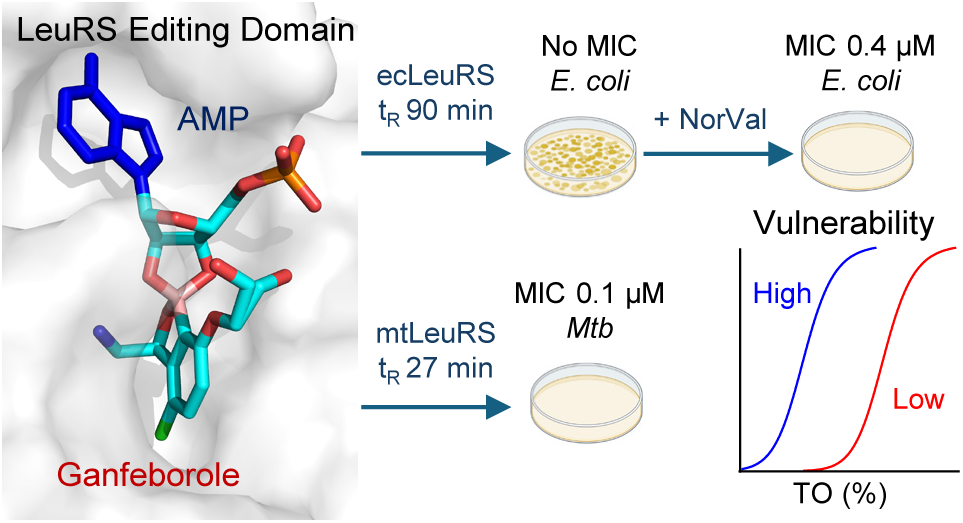

